# Galectin-3 inhibits *Paracoccidioides brasi liensis* growth and impacts paracoccidioidomycosis through multiple mechanisms

**DOI:** 10.1101/557918

**Authors:** Otavio Hatanaka, Caroline Patini Rezende, Pedro Moreno, Fabrício Freitas Fernandes, Patrícia Kellen Martins Oliveira Brito, Roberto Martinez, Carolina Coelho, Maria Cristina Roque-Barreira, Arturo Casadevall, Fausto Almeida

## Abstract

The thermodimorphic pathogenic fungi *Paracoccidioides brasiliensis* and *Paracoccioidioides lutzii* are the etiologic causes of paracoccidioidomycosis (PCM), the most prevalent systemic mycosis in Latin America. Galectin-3 (Gal-3), an animal β-galactoside-binding protein, modulates important roles during microbial infections, such as triggering a Th2-polarized immune response in PCM. Herein, we demonstrate that Gal-3 also plays other important roles in *P. brasiliensis* infection. We verified Gal-3 levels are upregulated in human and mice infections and establish that Gal-3 inhibits *P. brasiliensis* growth by inhibiting budding. Furthermore, Gal-3 affects disruption and internalization of extracellular vesicles (EV) from *P. brasiliensis* by macrophages. Our results suggest important roles for Gal-3 in *P. brasiliensis* infection, indicating that increased Gal-3 production during *P. brasiliensis* infection may account for affecting the fungal growth and EV stability, promoting a benefic course of experimental PCM.

**IMPORTANCE:** Paracoccidiodomycosis (PCM) is the most prevalent systemic mycosis in Latin America. Although the immune mechanisms to control PCM are still not fully understood, several events of the host innate and adaptive immunity are crucial to determine the progress of the infection. Mammalian β-galactoside-binding protein Galectin-3 (Gal-3) plays significant roles during microbial infections, and has been studied for its immunomodulatory roles but it can also have direct antimicrobial effects. We asked whether this protein plays a role in *P. brasiliensis*. We report herein that Gal-3 indeed has direct effects on fungal pathogen, inhibiting fungal growth and reducing extracellular vesicles stability. Our results suggest a direct role for Gal-3 in *P. brasiliensis* infection, with beneficial effects for the mammalian host.

## INTRODUCTION

Paracoccidioidomycosis (PCM), the most prevalent systemic mycosis in Latin America, is caused by the thermodimorphic human pathogens *Paracoccidioides brasiliensis* and *Paracoccioidioides lutzii*. After inhalation of airborne propagules from fungal mycelia phase, in the lungs the fungi convert into the infectious form - yeast phase (1–3). The yeast can spread to several organs causing systemic disease (4). Human defense against PCM depends on satisfactory cellular immune response and cytokine production (5, 6). Immune mechanisms that prevented cell division and budding of the fungal cells could aid in the control the PCM.

Extracellular vesicles (EVs) are produced by all living cells, and actively participate as key regulators of physiopathological mechanisms during fungal infections (7, 8). Fungal EVs carry several virulence factors and other important molecules, contributing to fungal pathogenicity and host immunomodulation (9–14). Since EVs plays significant roles in the host-pathogen relationship, the vesicular stability is important to ensure suitable delivery of their cargo into host cells (13, 15).

Bacterial and eukaryotic pathogens present surface glycans that may be recognized by host carbohydrate-binding proteins. These interactions commonly affect the microorganism pathogenesis, the host immune response or the success of intracellular parasitism (16–19). Recently, we have reported that Galectin-3 (Gal-3), a β-galactoside-binding animal lectin, plays significant roles in cryptococcal infection (13). Gal-3 interferes the *C. neoformans* infection, inhibiting *C. neoformans* growth and promoting vesicle disruption (13). Also, Gal-3 has been reported to influence the outcome of other mycoses, such as *Candida albicans* (20) and *Histoplasma capsulatum* (21). In *Paracoccidioides brasiliensis*, Gal-3 was reported to play an immunomodulatory role in the host response (22). Since Gal-3 can influence host response against PCM, as well as several other microbial infections, and regulates different functions in the physiopathology of infections, we explored whether Gal-3 influence *P. brasiliensis* growth and vesicle stability.

In this work, we assessed the Gal-3 levels in humans and mice with PCM. Also, we demonstrated the influence of Gal-3 in the *P. brasiliensis* growth and stability of EVs. Our results demonstrate that Gal-3 inhibits growth and budding of *P. brasiliensis* yeast cells, and promotes vesicle disruption. Our results suggest that Gal-3 can impacts the interaction of *P. brasiliensis* with host cells.

## RESULTS

### Gal-3 is up-regulated during PCM

Since increased Gal-3 expression was previously described during human and experimental inflammatory diseases (23, 24), and recently in *C. neoformans* infection (13), we measured Gal-3 levels in serum samples from individuals suffering of PCM, either acute or chronic PCM form. Compared to the healthy individuals, the acute and chronic form patients showed higher Gal-3 serum levels, as shown previously for other infections (Figure 1). There was no significant difference (*P* value: 0.4204, unpaired t-test) between the Gal-3 levels in sera of acute and chronic patients with PCM.

**Figure 1.**
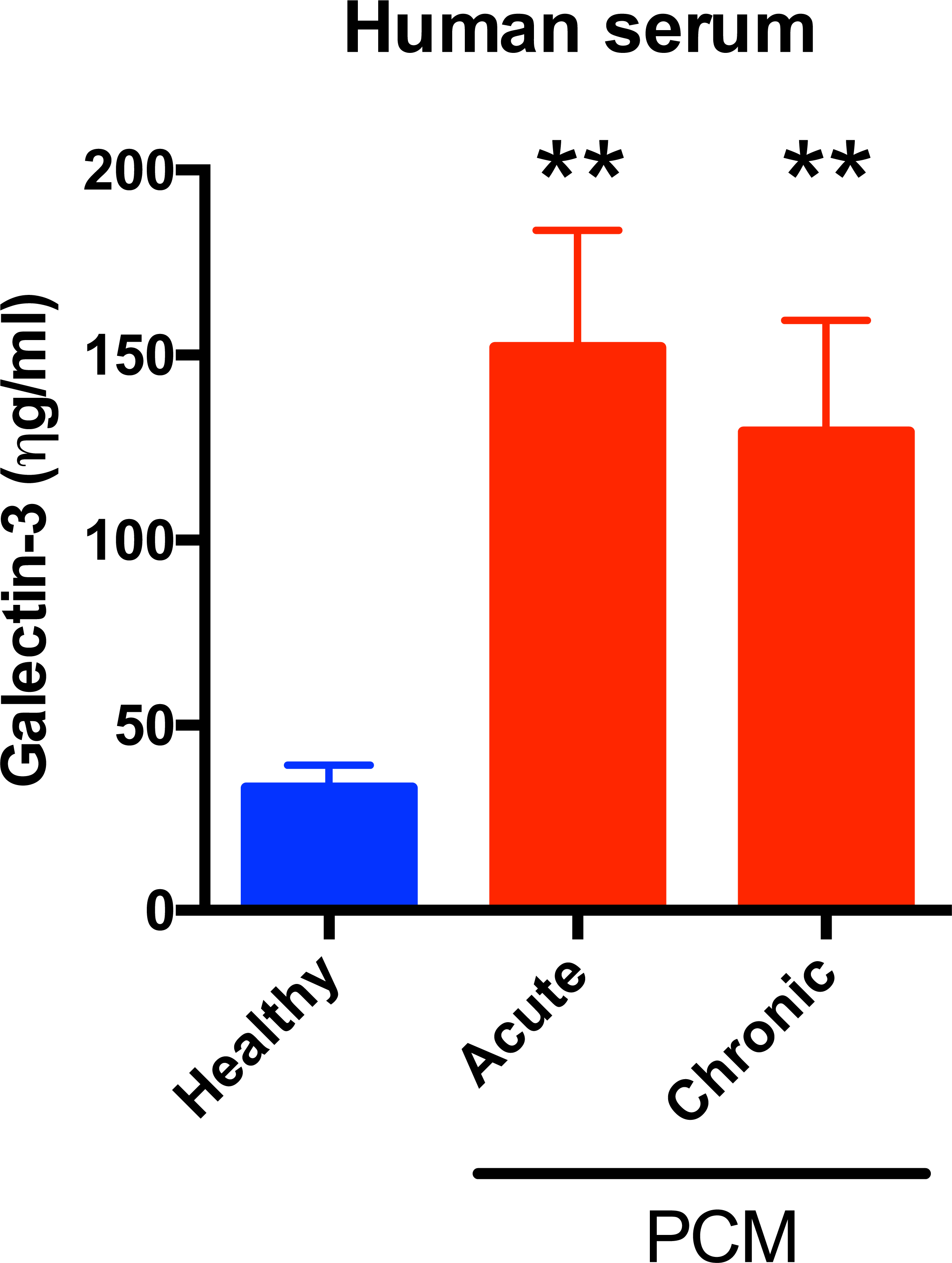
Upregulated Gal-3 levels in humans during *P. brasiliensis* infection. Gal-3 levels in serum from healthy individuals (blue bar) and patients infected by *P. brasiliensis* (red bars) were assessed by ELISA. Gal-3 levels were higher in acute and chronic form patients infected with *P. brasiliensis* when compared with healthy individuals. Bars represent the mean ± SD of Gal-3 levels obtained from triplicate samples. Statistically significant differences are denoted by asterisks (** p < 0.005, unpaired Student’s t-test).

Subsequently, we measured Gal-3 levels in tissues and serum of C57BL/6 mice on days 30 and 60 post-infection with *P. brasiliensis* (Figure 2). In comparison with control animals (PBS), infected mice had higher Gal-3 levels in all examined tissues (lungs and spleen, Figure 2B and 2C, respectively) and serum samples (Figure 2A). As previously reported for *C. neoformans* infection, there is a correlation between *P. brasiliensis* infection and increased levels of serum Gal-3, which could reflect the inflammatory conditions caused by these infectious diseases.

**Figure 2.**
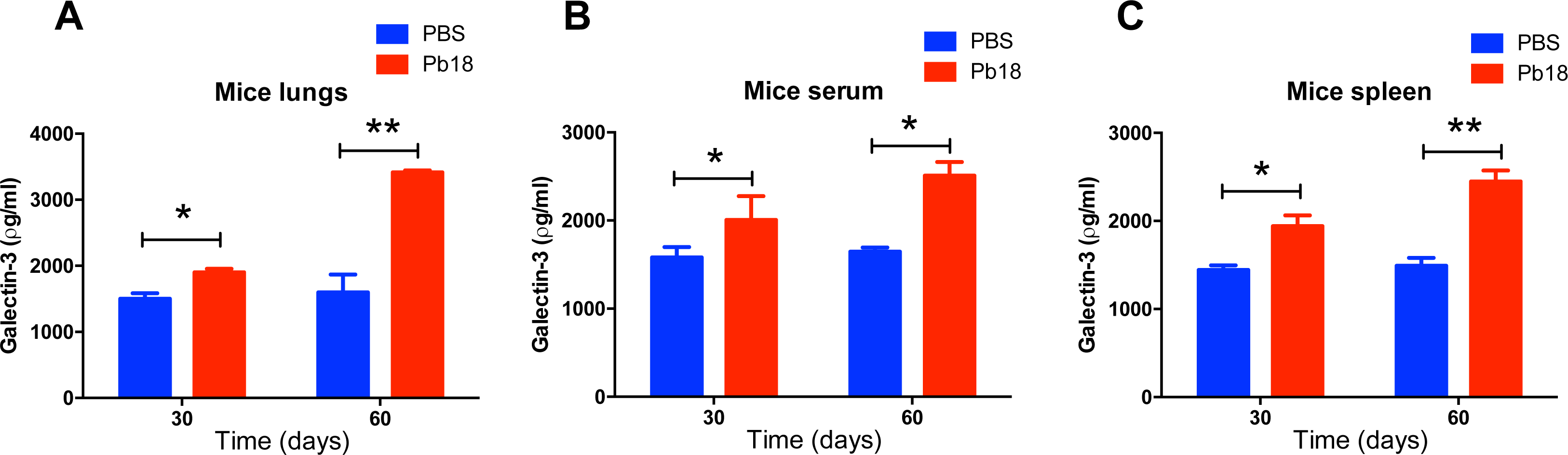
Upregulated Gal-3 levels in mice during experimental *P. brasiliensis* infection. C57BL/6 mice were intratracheally infected with Pb18 strain yeast cells (red bars) or PBS (blue bars) and Gal-3 levels were assessed in tissues and serum during the course of *P. brasiliensis* infection. On days 30 and 60 after infection, samples collected of lung (A), serum (B), and spleen (C) were homogenized and Gal-3 quantified by ELISA. Bars represent the mean ± SD of Gal-3 levels obtained from triplicate measurements for each animal, with five animals per group. Statistically significant differences are denoted by asterisks (* p < 0.05, ** p < 0.005, unpaired Student’s t-test).

### Gal-3 inhibits *P. brasiliensis* growth

Since *P. brasiliensis* cell division and budding is crucial to successful PCM, and Gal-3 inhibits *C. neoformans* growth (13), we evaluated whether Gal-3 could affect the growth and budding of *P. brasiliensis*. *P. brasiliensis* growth in culture, measured by MTT assay, was compared between Gal-3 treated, PBS-and denatured Gal-3-treated yeasts. Gal-3 inhibited *P. brasiliensis* growth by approximately 50% after 72 h compared with the controls (denatured Gal-3 treated or PBS) (Figure 3). To verify whether Gal-3 treatment of fungal yeast induces yeast death, we performed viability assays using fluorescein diacetate/ethidium bromide staining. Gal-3 treated and control cultures contained similar proportions of viable cells up until 72 h after Gal-3 treatment, and all cultures were >80% viable (data not shown). To further characterize Gal-3 effects in the growth of *P. brasiliensis*, we evaluated the average number of cells with buds in the yeast culture in the presence or absence of Gal-3 for 72h, as well as in the presence of denatured Gal-3. We counted the budded or unbudded yeast cell via direct observation in a Neubauer chamber (Figure 3B). The average number of budding cells was 80% in both untreated and denatured Gal-3-treated cells. On the other hand, the average was decreased to 49% in Gal-3-treated cells.

**Figure 3.**
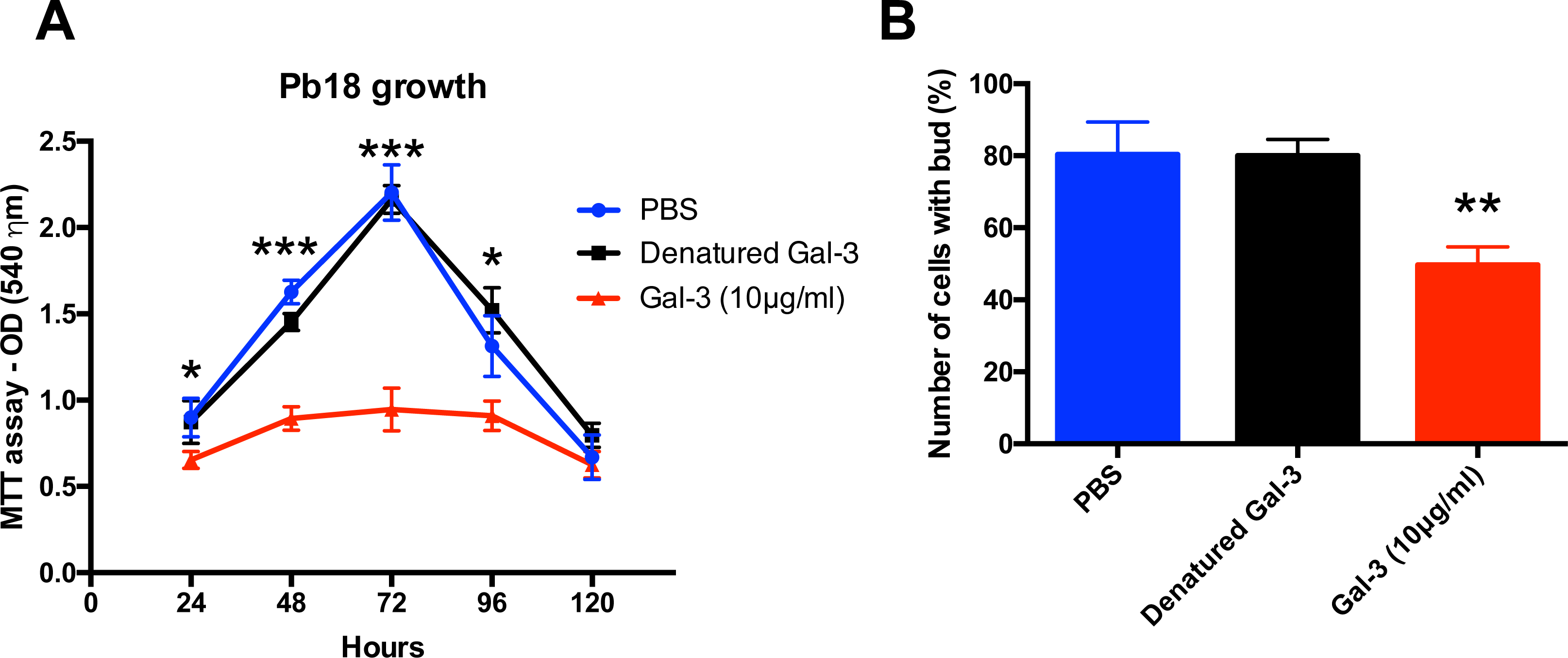
Gal-3 inhibits the growth and budding of *P. brasiliensis* yeast cells. MTT assay of *P. brasiliensis* Pb18 strain cultivated in YPD medium for 120 hours at 37°C with 10 µg/ml of Gal-3 or denatured 10 µg/ml Gal-3 (A). Number of cells with buds in YPD medium in the absence (blue bar), or presence of denatured Gal-3 (black bar), or presence of 10 µg/ml Gal-3 (red bar) for 72 hours at 37°C. Buds were counted via light microscopy and quantified using a Neubauer chamber hemocytometer. (B). Data are representative of three experiments showing mean ± SD for each data point. Statistically significant differences are denoted by asterisks (* p < 0.05, ** p < 0.005, *** p < 0.0005, unpaired Student’s t-test).

Flow cytometry assessment of the Gal-3 binding to *P. brasiliensis* Pb18 strain showed that Gal-3 bound to *P. brasiliensis* cells (Figure 4A). Confocal microscopy demonstrated that Gal-3 co-localized with calcofluor white, a cell wall dye (Figure 4B-D and Supplementary Figure 1). Calcofluor white was used as a positive control as it binds to the cell wall (CW). These results suggest that the recognition of the fungal cell wall by Gal-3, through an unknown sugar moiety, may explain its inhibitory effect in *P. brasiliensis in vitro* growth.

**Figure 4.**
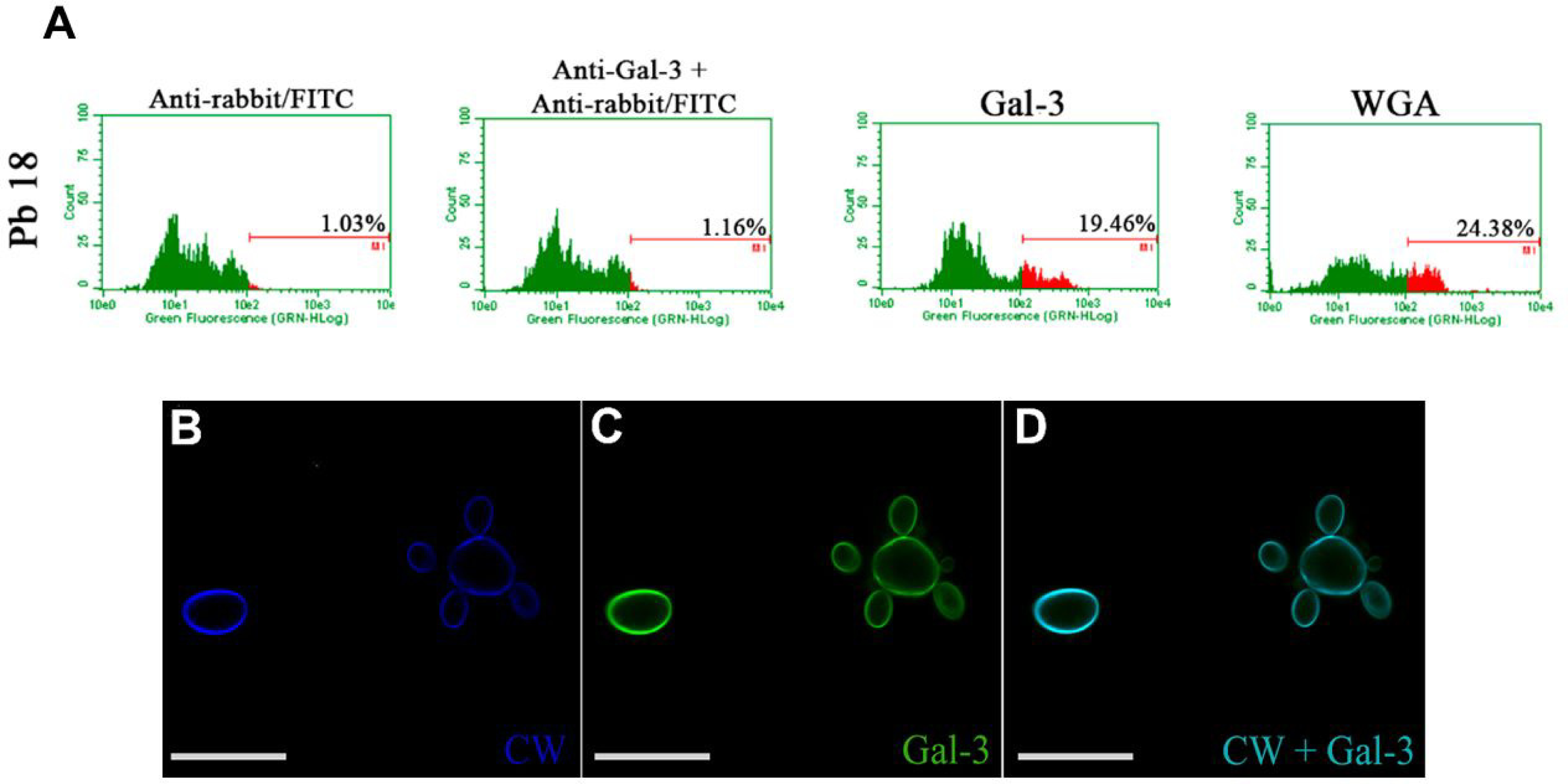
Gal-3 binds to *P. brasiliensis* cell wall. *P. brasiliensis* Pb18 strain cultivated in YPD medium for 72 hours at 37°C, was resuspended in PBS and incubated sequentially with 40 µg/ml of Gal-3, an anti-Gal-3 antibody and finally anti-rabbit IgG-FITC antibody. Binding was measured by flow cytometry; numbers inside histogram represents the percentage of positive cells recognized by Gal-3 (A). WGA lectin-FITC (30 μg/ml) as control of binding with cell wall (CW) (A). *P. brasiliensis* Pb18 strain was cultured at 37°C for 72 h and incubated with Gal-3. *P. brasiliensis* were stained for observation of cell wall with calcofluor white (blue) (B) and Gal-3 with anti-Gal-3 antibody (green) (C). Merged images are shown in D (cell wall and Gal-3). The images represent a single section from a Z series stack. Scale bar correspond to 10 µm. Data are representative of three experiments and a representative image is shown.

### Gal-3 disrupted *P. brasiliensis* EVs

Exposure of EVs produced by *C. neoformans* to Gal-3, macrophages, or bovine serum albumin causes vesicular disruption (13, 15). We asked whether Gal-3 would affect the stability of EVs produced by *P. brasiliensis*. Addition of Gal-3 to radiolabeled EVs promoted vesicular disruption and subsequent radioactive release in a dose-dependent manner (Figure 5A). Furthermore, we blocked Gal-3 carbohydrate recognition domain (CRD) by pre-incubating with N-acetyl-lactosamine (lactosamine, Gal-3 glycoligand) and by Gal-3 denaturation (boiling at 10 min for 100 °C). Both denatured and lactosamine-bound Gal3 had no lytic effect on *P. brasiliensis* EVs (Figure 5A), suggesting that intact 3D conformation and Gal-3 CRD are important for the Gal-3 lysing activity. Moreover, radioactive assays showed that other lectins were unable to lyse *P. brasiliensis* EVs (Figure 5B).

**Figure 5.**
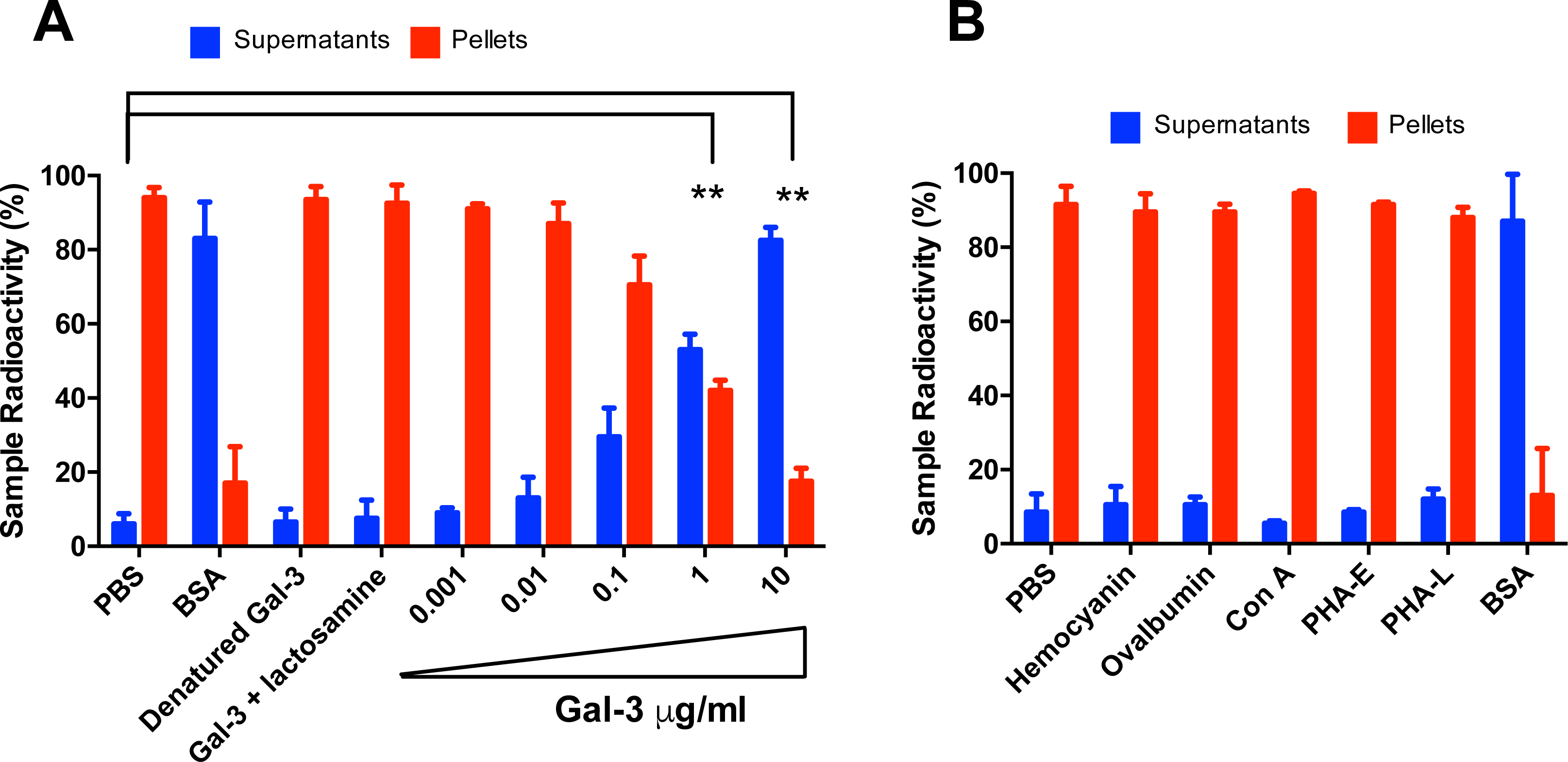
Gal-3 disrupts *P. brasiliensis* extracellular vesicles. Purified radiolabelled vesicles after 72 h post [1-^14^C] palmitic acid addition were resuspended in PBS, BSA, denatured Gal-3, Gal-3 pre-incubated with lactosamine, Gal-3 (0.001 to 10 µg/mL) and release of radioactivity from vesicle pellet was assayed. Supernatant (blue bars) and pellet (red bars) radioactivity were assessed and normalized to 100% radioactivity for each individual sample (A). Hemocyanin, ovalbumin, concanavalin A (Con A), phytohaemagglutinin E (PHA-E), phytohaemagglutinin L (PHA-L), and bovine serum albumin (BSA) were used as control (B). Bars represent the mean ± SD from triplicate samples from one representative experiment. Experiments were repeated at least xx times. Statistically significant differences are denoted by asterisks (** p < 0.005, unpaired Student’s t-test).

### Gal-3 affects macrophages capability to disrupt and internalize EVs

Given that Gal-3 is expressed and play a myriad roles in macrophage populations (25–27) and on the previous observation that macrophages (15) and Gal-3 can disrupt EVs from *C. neoformans* (13) and *P. brasiliensis* (this study), we evaluated whether these Gal3 binding and EV lysis could be correlated for *P. brasiliensis*. The addition of radiolabeled EVs from *P. brasiliensis* to the macrophages showed that WT peritoneal macrophages were approximately three times more effective than *Gal-3^−/−^* macrophages to disrupt EVs (black bars, Figure 6). Moreover, we demonstrated that uptake of EVs from *P. brasiliensis* by WT peritoneal macrophages gradually increased, whereas the uptake by *Gal-3^−/−^* peritoneal macrophages increased significantly less (Figure 6).

**Figure 6.**
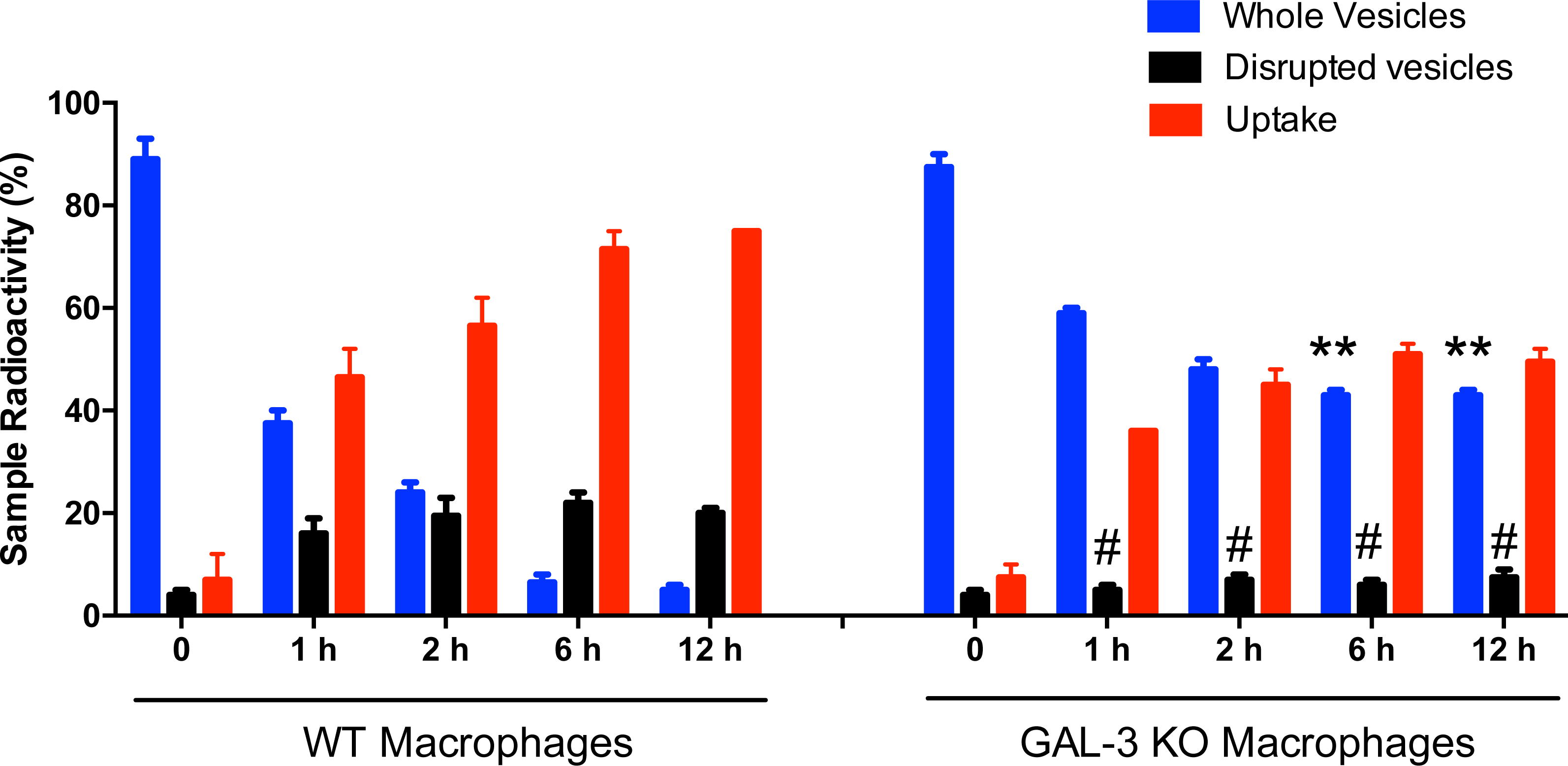
Gal-3 affects disruption and internalization of *P. brasiliensis* EVs by macrophages. Purified radiolabelled EVs from *P. brasiliensis* Pb18 strain yeast cultures were added to cultures of C57BL6 WT or Gal-3^−/−^ macrophages. After 1, 2, 6 or 12 h post EVs addition, the radioactivity recovered from the macrophages (adhered cells, uptake, red bars), whole vesicles (pellet, blue bars) and disrupted vesicles (supernatant, black bars) were quantified. Bars represent the mean ± SD from triplicate samples from 3 representative experiments. Statistically significant differences are denoted by asterisks (** p < 0.005, unpaired Student’s t-test).

## DISCUSSION

Herein we describe a role for Gal-3 in *P. brasiliensis* infection that parallels recent observations with *C. neoformans* (13). Galectins are able to regulate positively or negatively host-microbial interactions in respiratory infections according to galectin type, pathogen and host context (28, 29). Gal-3, member of the galectin family of β-galactosides-binding proteins, is widely expressed in different cells, and plays important roles in biological phenomena, such as inflammation and immunity (28, 30).

Our previous results in experimental models of cryptococcosis showed Gal-3 were increased during infection (13). This observation was replicated in human serum where higher levels of Gal-3 were detected in patients with cryptococcosis when compared with healthy individuals (13). The low number of patients available for study prevented us from making definitive statements regarding increase of Gal-3 in human infection, but in a prior study (13) Gal-3 was shown to be increased in infection and therefore it is very likely that the increase we observed is true in *P. brasiliensis* infection. Moreover, our results show that Gal-3 inhibits the fungal growth and morphogenesis of *P. brasiliensis,* a fungistatic effect of Gal-3 comparable to what was previously verified for *C. neoformans* (13). In mouse models deficiency of Gal-3 lead to an increased microbe burden and a decreased survival of the animals (13, 20, 22, 31). Furthermore *gal3^−/−^* mice die faster than wild-type mice when infected with *P. brasiliensis* (22). Gal-3 antifungal effects are widely conserved, affecting most (if not all) fungal pathogens and reaffirming Gal-3 as a critical player in antifungal defenses.

Gal-3 promoted disruption of *C. neoformans* EVs and influenced the uptake of EV content by macrophages (13), and we now replicate these observations for *P. brasiliensis*. The disruption mechanism of EVs remains cryptic. As discussed previously (13), albumin induce vesicles disruption (15) and albumin can bind to fatty acids (32) and sterols (33) and may promote membrane destabilization. However we cannot propose a similar mechanism for Gal-3 mediated disruption of EVs, particularly since Gal-3 is not known to bind lipids. We showed Gal-3 disrupts EVs in a manner dependent on denaturation and the CRD domain, and we hypothesize the existence of a novel mechanism of EV disruption. Gal-3 binds beta-galactosides, in proteins (34) and microbe surfaces (specifically the fungal cell wall), and it is likely that a version of this galactoside is displayed on the surface of microbial EVs. Further studies are needed to discern the glycan moiety recognized in fungal EVs (and whether the glycan moiety is associated to proteins or instead to an putative carbohydrate polymers present in EVs) and how this binding triggers collapse of the lipid bilayer to disrupt EVs. In any case, this may constitute an important mechanism of immune defense: Gal-3 lysis of EVs would prevent EVs’ delivery of a concentrated cargo of virulence factors at the host cell surface and instead result in diluted release of fungal components into the extracellular milieu and heightened degradation by host extracellular enzymes.

We conclude that Gal-3 is beneficial for the mammalian host during *P. brasiliensis* infection by contributing to host defense. As previously reported for *C. neoformans* (13), the antimicrobial mechanism of Gal-3 is due to a combination of microbial vesicle lysis, coupled to inhibition of fungal growth and morphogenesis. In addition to direct antimicrobial effects, Gal-3 plays immunomodulatory roles (31, 35–37) that may synergize with the antimicrobial effects. Multiple inhibitors of Gal-3 have been designed (38) – and are being tested for antitumorigenic properties. However our data reveals that these therapies may predispose patients for fungal infections and, as is the case for many immunotherapies, it is important to closely monitor fungal infections in these patients. In the case of fungal infections, it may be desirable to design a Gal-3 mimetic that, through inhibition of growth and interference with EV release, would act as a potent antifungal therapy.

## METHODS

### Ethics statement

All animal use complied with the standards described in Ethical Principles Guide in Animal Research adopted by the Brazilian College of Animal Experimentation. The protocols were approved by the Committee of Ethics in Animal Research of the Ribeirao Preto Medical School at the University of Sao Paulo (protocol 20/2013-1). Informed written consent from all participants was obtained. The studies involving patients were approved by the Research Ethics Committee of the University Hospital, Ribeirao Preto Medical School at the University of Sao Paulo (protocol HCRP 13.982/2005).

### Mice and *P. brasiliensis* strain

We used male C57BL/6 (wild-type, WT, Jax 000664) and galectin-3-deficient mice (*gal3^−/−^*) at 6 to 8 weeks of age. Knockout mice were kindly donated by FT Liu (University of California, Davis, CA). *Gal3^−/−^* mice were previously generated as described and crossbred to the C57BL/6 mouse background for nine generations (39). The animals were housed in the animal facility of the Ribeirao Preto Medical School, University of São Paulo, under optimized hygienic conditions. All *P. brasiliensis* experiments were performed with the Pb18 isolate. Fungal cultures were grown in the YPD medium (2% peptone, 1% yeast extract, and 2% glucose) in the yeast phase, at 36 °C. To ensure yeast virulence, serial passages in BALB/c mice were performed before the isolate Pb18 was used in experiments.

### Sera and patients

Blood samples were obtained from patients being seen in the University Hospital, Ribeirao Preto Medical School at the University of Sao Paulo. A total of 6 patients with diagnosed paracoccidioidomycosis were included in this study: 3 patients with acute and 3 patients with chronic form. Serum was obtained and stored at −80°C. Samples were also obtained from 3 blood donors with a median age of 30 years (range, 25-35 years).

### Gal-3 levels

Gal-3 levels in the lung, serum and spleen was quantified in the organ homogenates of *P. brasiliensis*-infected mice. The homogenate samples (whole organ in 1 ml of PBS) of control mice or from animals infected with *P. brasiliensis*, as well as the serum samples from infected animals or from patients with diagnosed paracoccidioidomycosis, were stored at −80°C until assayed. All samples were thawed only once prior to use. Gal-3 levels were measured using commercially available enzyme-linked immunosorbent assay (ELISA) kits (Sigma-Aldrich, St. Louis, MO, USA) according to the manufacturer’s instructions.

### Cell viability and growth in the presence of Gal-3

Fungal viability was determined using fluorescein diacetate/ethidium bromide staining. Only the cultures that were greater than 85% viable were used.

To verify Gal-3 effect on the cells, we performed growth curves in YPD liquid medium containing different concentrations of Gal-3 (Gal-3 human recombinant, expressed in *E. coli*, Sigma-Aldrich) in a 96-well plate (Costar, NY, USA). The Gal-3 effect on cell proliferation was determined using an MTT assay (40), as follows: *P. brasiliensis* yeast cells were suspended in YPD medium at a density of 10^6^ cells/ml, and Gal-3, denatured Gal-3 or PBS were used as control. After incubation at 37°C in an orbital shaker (150 rpm) for 120 h. To verify the Gal-3 effect in the yeast budding, we assessed the average number of Pb18 strain cells with buds found in yeasts cultures after 72 h of Gal-3 treatment and compared to the cells treated with denatured Gal-3 and PBS treated. Counting was carried out in a neubauer chamber by optical microscopy, considering as budding cells, the yeasts that presented at least one or more bud.

### Gal-3 binding assay and confocal microscopy

The Gal-3 binding assay and confocal microscopy were performed as previously described for *C. neoformans* (13) using *P. brasiliensis* yeast cells. Gal-3 binding assay was performed with yeast cells incubated with PBS containing 10% fetal bovine serum for 20 min at 4°C to block non-specific binding of antibodies. Next, 1 ml of the suspension containing 10^6^ cells were incubated with either Gal-3 (40 µg/ml) and denatured Gal-3 (40 µg/ml) for 40 min at 4°C. Cells were washed twice with PBS and anti-Gal-3 antibody (1:50; Sigma-Aldrich) was added; after incubation for 45 min, the cells were washed twice with PBS and incubated with anti-rabbit IgG-FITC antibody (1:50; Sigma-Aldrich) for 40 min at 4°C. Gal-3 binding to *P. brasiliensis* cells was analyzed by flow cytometry (Guava easyCyte, Guava Technologies, Millipore, Hayward, CA, USA). Anti-rabbit IgG-FITC antibody associated or not with Gal-3 and WGA lectin (30 µg/ml) were, used as negative and positive controls, respectively. The confocal microscopy was performed with cells incubated with Gal-3 (40 µg/ml) at 37°C for 1 h followed by three washes with PBS and fixation with PBS-buffered 3.7% formaldehyde at 25°C. The samples were washed three times with PBS and treated with glycine 0.1 M for 15 min, and blocked with BSA (1% in PBS) for 1 h at 25°C. Then, the cells were incubated with a rabbit anti-Gal-3 antibody (Sigma-Aldrich) overnight at 4°C. The samples were washed five times with PBS and incubated for 1 h with FITC-labeled donkey anti-rabbit IgG from Jackson Immuno Research Laboratories. For cell wall staining, samples were incubated with Calcofluor White (50 µg/ml, Sigma-Aldrich) in PBS for 20 min. After five washes with PBS, cells were placed on slide and coverslips mounted with Fluoromount-G (Electron Microscopy Sciences). The samples were examined with a LSM780 system AxioObserver, 63X oil immersion (Carl Zeiss, Jena, Germany). The images were analyzed offline using the ImageJ software (http://rsb.info.nih.gov/ij/). Secondary antibody alone was used as controls. All controls were negative.

### Analysis of the stability of extracellular vesicles

EVs were isolated as previously described (41). Vesicles quantification was measured by Nanoparticle-Tracking Analysis (NTA) using a NanoSight NS300 (Malvern Instruments, Malvern, UK) equipped with fast video capture and particle-tracking software, as previously described (14). Purified vesicles from *P. brasiliensis* were diluted into PBS, and each sample was then injected into a NanoSight sample cubicle. The measurements were obtained in triplicate and analyzed using NanoSight software (version 3.2.16). The EVs stability was evaluated according to protocols previously described (13). EVs were incubated with Gal-3 (Gal-3 human recombinant, expressed in *E. coli*, Sigma-Aldrich) at different final concentration (0 to 10 µg/ml), and the concentrations of all control lectins were normalized according to carbohydrate binding sites. EVs stability was examined by radioactive assay through cultivation of *P. brasiliensis* in the presence of [1-^14^C] palmitic acid, as previously described for *C. neoformans* (13, 15). The suspension of radiolabeled EVs was incubated with Gal-3 at 37°C for different times and concentrations, and the suspension was ultracentrifuged at 100,000 × g for 1 h at 4°C. Supernatants and pellets were saved for scintillation counting.

### Vesicle disruption and uptake by macrophages

To assess the vesicle stability and vesicle uptake by macrophages from WT and gal3^−/−^ mice, we used the protocol previously described (13). Peritoneal macrophages were obtained from C57BL/6 WT or gal3^−/−^ mice, and grown in DME medium (Invitrogen) supplemented with 10% (v/v) fetal bovine serum, 10% NCTC (Invitrogen), 1% nonessential amino acids (Invitrogen) and 1% penicillin (Invitrogen). 48-well tissue culture plates were seeded with elicited peritoneal macrophages (4 × 10^5^ cells/well). EVs were obtained from *P. brasiliensis* cultures that were pulsed with [1-^14^C] palmitic acid 72 h before EVs harvesting, and added to macrophage cultures as previously described (13). The adhered cells (containing EVs due to uptake), supernatants (containing components of disrupted EVs) and pellets (containing intact EVs) were saved for scintillation counting. The radioactivity distribution in the three fractions was expressed as percent of the total radioactivity.

### Statistical Analysis

Data are either the means of or representative results from at least 3 independent experiments, each performed in triplicated. All statistical analyses and comparisons were performed using the *GraphPad Prism* Software version 6.0 (GraphPad Software, San Diego, CA, USA). A *P* value < 0.05 was considered statistically significant.

## ACKNOWLEDGEMENTS

We thank Patricia Vendruscolo and Roberta Ribeiro Costa Rosales from Ribeirao Preto Medical School, Sao Paulo, Brazil, for technical support. FA received funding from Fundação de Amparo à Pesquisa do Estado de São Paulo (2016/03322-7, 2016/15055-3) - Project Young Researcher, CNPq (Conselho Nacional de Desenvolvimento Científico e Tecnológico, and CAPES (Coordenação de Aperfeiçoamento de Nível Superior). AC was supported in part by NIH awards AI033142, AI052733 and HL059842.

## AUTHORS CONTRIBUTIONS

All of the authors contributed to the research design and data analyses. Performed the experiments: OH, CP, PM, FF, PK, FA. Contributed reagents/materials/analysis tools: RM, MC, AC, FA. Wrote the paper: OH, CP, PM, CC, MC, AC, FA.

## COMPETING FINANCIAL INTERESTS

The authors declare no competing financial interests.

---

**Supplementary Figure 1. Co-localization of Gal-3 and cell wall of *P. brasiliensis*.** Co-localization of Gal-3 binding sites with cell wall structures was confirmed by the threshold Mander’s coefficient tool available in the Fiji J software. Analysis by fluorescence microscopy included cell wall (Calcofluor white, CW, blue fluorescence) and Gal-3 binding sites (anti-Gal-3 antibody, green fluorescence). The whole image field was used to obtain the split threshold Mander’s colocalization coefficient for each channel (Tm1/Tm2).

